# The Retrosplenial Cortical Role in Delayed Spatial Alternation

**DOI:** 10.1101/2024.06.18.599656

**Authors:** Dev Laxman Subramanian, Adam M.P. Miller, David M. Smith

## Abstract

The retrosplenial cortex (RSC) plays an important role in spatial cognition. RSC neurons exhibit a variety of spatial firing patterns and lesion studies have found that the RSC is necessary for spatial working memory tasks. However, little is known about how RSC neurons might encode spatial memory during a delay period. In the present study, we trained control rats and rats with excitotoxic lesions of the RSC on spatial alternation task with varying delay durations and in a separate group of rats, we recorded RSC neuronal activity as the rats performed the alternation task. We found that RSC lesions significantly impaired alternation performance, particularly at the longest delay duration. We also found that RSC neurons exhibited reliably different firing patterns throughout the delay periods preceding left and right trials, consistent with a working memory signal. These differential firing patterns were absent during the delay periods preceding errors. We also found that many RSC neurons exhibit a large spike in firing rate leading up to the start of the trial. Many of these trial start responses also differentiated left and right trials, suggesting that they could play a role in priming the ‘go left’ or ‘go right’ behavioral responses. Our results suggest that these firing patterns represent critical memory information that underlies the RSC role in spatial working memory.

## 1. Introduction

The retrosplenial cortex (RSC) is a key component of the memory and navigation systems of the brain (for reviews see, Mitchell et al., 2018, Smith et al., 2022, Vann et al., 2009). Previous studies have found that RSC lesions impair performance on a variety of spatial navigation tasks (Parron & Save, 2004; Pothuizen et al., 2008; Sutherland et al., 1988; Vann & Aggleton, 2002). The RSC may be particularly important for spatial working memory. For example, RSC lesions impair performance on the eight-arm radial maze (Keene & Bucci, 2009). However, the effects of RSC lesions can be subtle and the lesion-induced deficits are variable, depending on the specific experimental procedures (Hindley et al., 2014; Nelson et al., 2015; Vann et al., 2003, Vann & Aggleton, 2005). RSC lesion-induced impairments are delay-dependent, with more severe impairments at longer delays (Keene & Bucci, 2009). The anatomical extent of the lesions within the RSC is also important, since partial lesions produce a milder impairment than more complete lesions (Vann & Aggleton, 2004). Consistent with this, we examined the role of the RSC in a continuous (non-delayed) version of the T-maze alternation task and found that the average lesion-induced impairment was mild. However, the impairment was tightly correlated with the size of the lesions (Miller et al., 2019). In the present study, we examine the effects of RSC lesions on T-maze alternation with a variable delay.

A growing literature shows that RSC neurons exhibit a wide range of navigation-related firing patterns in rodents (e.g. Subramanian et al., 2024). These firing patterns carry information about the subject’s spatial location (Miller et al., 2019), head direction (Cho & Sharp, 2001; Jacob et al., 2017), routes and trajectories (Alexander & Nitz, 2015, 2017), environmental boundaries (Alexander et al., 2020), navigational cues (Vedder et al., 2017), and goal locations (Miller et al., 2019). Our previous study of continuous alternation (Miller et al., 2019) found that RSC firing patterns differentiated the left and right trials and simulated the upcoming goal locations as the rats traversed the stem of the maze, suggesting that these neurons carry important memory information about the reward locations. Less is known about delay-related firing patterns in the RSC. In a virtual navigation task, context-specific firing patterns during the trials persisted into the subsequent delay period (Sun et al., 2021), but it is not clear whether these firing patterns were relevant to memory. More recently, we found that RSC neurons exhibit time fields, characterized by discrete periods of activity during specific epoch of the delay period (Subramanian & Smith, 2024). Notably, these firing patterns were sensitive to the memory requirements of the task. In the present study, we examined RSC firing patterns during the delay period of the T-maze alternation task to determine whether they could support working memory representations.

## 2. Methods

### 2.1 Subjects

Subjects were 28 adult male Long-Evans rats obtained from Charles River Laboratories, Wilmington, MA weighing 250g-300g upon arrival. Eight rats were used in the electrophysiology study and 20 rats were used in the lesion study. Of the 10 rats that received RSC lesions, 2 were excluded from the analysis due to hippocampal damage, and 1 was excluded because the RSC damage was unilateral. Rats were placed on a 12hr/12hr light/dark cycle with lights on at 8am and allowed to acclimate to the vivarium for at least one week prior to surgery. After surgery, rats were placed on food restriction until they reached 80-85% of their free-feeding weight. Water was always available ad libitum. All procedures complied with the guidelines established by the Cornell University Animal Care and Use Committee.

### 2.2 Surgical procedures

#### 2.2.1 Lesions

The rats were anesthetized with isoflurane and placed in a stereotaxic apparatus. The skin was retracted, and holes were drilled through the skull above each of the injection sites. Rats assigned to the lesion condition were given bilateral injections of N methyl-D-aspartate (NMDA, 10μg/ml) in 6 injection sites per hemisphere, with stereotaxic coordinates (Paxinos & Watson, 1998) and injection volumes as follows: 0.35μL at −2.2 (AP), ±0.5 (ML), −3.0 (DV); 0.35μL at −3.9, ±0.5, −3.0; 0.20μL at −5.5, ±0.5, −3.5; 0.35μL at −5.5, ±1.0, −2.8; 0.35μL at −6.7, ±1.1, −2.8; 0.30μL at −8.0, ±1.3, −2.8. Coordinates were taken from Bregma (AP), the midline (ML), and the surface of the skull (DV), respectively. The injection cannula was left in place 1 min before and 5 min after each infusion. Control rats were given sham lesions of the RSC consisting of lowering the injection cannula into the brain but not injecting NMDA.

#### 2.2.2 Electrode Microdrives

Eight rats had a custom-built electrode microdrive implanted, which contained 20 moveable tetrodes (16 recording tetrodes and 4 reference electrodes) made from twisting four 17μm platinum/iridium (90%/10%) wires, platinum plated to an impedance of 100-500kΩ, and arranged in two linear arrays of 10 tetrodes (one in each hemisphere) that spanned approximately 5mm along the rostro-caudal axis of the brain. Tetrodes were stereotaxically positioned bilaterally just beneath the cortical surface (2-7mm posterior to Bregma, ±1.5mm lateral) with the tetrodes angled 30 degrees toward the midline. Rats were given 7 days to recover from surgery prior to lowering the tetrodes into the RSC (35-70μm daily) over the course of several days until a depth of at least 1mm was reached in order to target the granular b subregion.

### 2.3 Behavioral procedure

The rats were trained on a black PVC continuous T-maze (120 cm long stem x 100 cm wide x 68 cm above the floor, Fig. 1A) with metal reward cups embedded in the ends of the arms through which chocolate milk (0.2 ml, Nestle’s Nesquik) was delivered via an elevated reservoir controlled by solenoid valves activated by foot-pedal switches. The maze was located in the center of a circular arena enclosed by black curtains with distinctive visual cues. A continuous background masking noise (white noise) was played from a speaker located directly above the maze. After acclimation to the maze, rats were trained on a continuous spatial alternation task in which they were rewarded only if they approached the reward location (left or right) which was opposite from the previous trial. Both cups were baited and the rats could choose either reward location on the first trial. Entries into the same arm as the previous trial were scored as an error and were not rewarded. Rats were not allowed to correct their errors and were instead gently ushered back if they left the continuous alternation route. Rats were given 40 trials/day until they achieved a criterion of 90% correct on two consecutive sessions.

**Figure 1.**
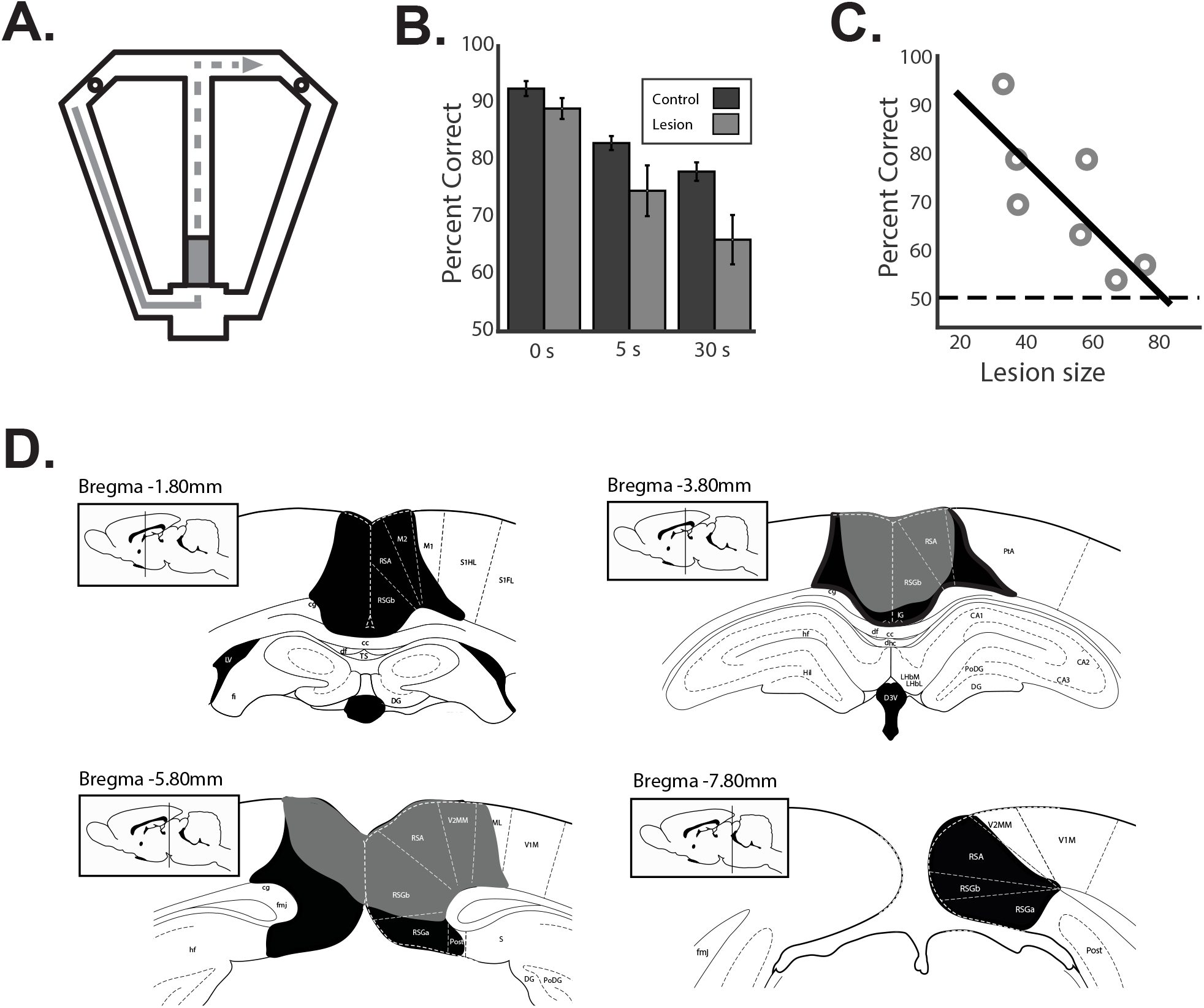
RSC lesions impair spatial working memory. A) illustrates the delayed alternation task. After visiting one of the two reward locations (circles), rats returned to the stem and had to approach the opposite location for reward on the following trial. A wooden block (grey rectangle) prevented the rat from entering the stem for a delay period of 0, 5 or 30 sec. B) Behavioral performance of control rats and rats with RSC lesions for the three delay durations. C) Performance at the 30-sec delay was strongly correlated with the extent of the RSC lesions. Chance performance is indicated by the dashed line. D) The largest (black) and smallest (grey) lesions are shown at four AP coordinates.

After achieving this criterion, rats were then trained on a delayed version of the task in which the path to the reward was blocked by a black wooden barrier placed at the entry to the stem of the maze. In the lesion experiment, we tested performance under three different delay conditions, 0 sec (block was removed as soon as the rat approached), 5 sec, or 30 sec. The 0 sec delay condition served to control for disruption of continuous locomotion or potential distraction caused by placement or removal of the block. In the electrophysiology experiment, all of the trials involved a 30 sec delay. The experimenter monitored delay lengths with a stopwatch.

### 2.4 Recordings

Neuronal spike data and video recordings of the rat’s location and head direction were collected using the Cheetah Digital Data Acquisition System (Neuralynx, Inc. Bozeman, MT). Neuronal recordings were filtered at 600Hz and 6kHz, digitized and stored to disc along with timestamps for offline sorting (SpikeSort3D, Neuralynx, Inc.). The rat’s position and head direction were monitored by digitized video (sampled at 30 Hz) of two LEDs attached to the head stage. The time of reward receipt was measured with a grounding circuit that detected oral contact with the chocolate milk reward. A total of 144 RSC neurons were recorded from 8 rats while they performed the delayed spatial alternation task.

### 2.5 Histology

After completion of the experiment, rats were transcardially perfused with 4% paraformaldehyde in phosphate buffered saline. Brains were removed and stored for at least 24hrs in 4% paraformaldehyde before being transferred to a 30% sucrose solution for storage until slicing. Coronal sections (40μm) were stained with 0.5% cresyl violet for visualization of tissue damage (in the case of NMDA lesions) or tetrode tracks (for neuronal recordings). Tissue damage was quantified by laying a grid (250μm to-scale grid spacing) over an enlarged image of the stained tissue and dividing the number of grid intersections located over damaged RSC areas by the number of intersections located over the entire RSC (Paxinos & Watson, 1998). Tetrode positions were identified using depth records noted during tetrode lowering and tracks observed in the stained tissue. Neuronal records from tetrodes located outside of the RSC were excluded from the data set. Our recordings targeted the granular b subregion of the RSC, although small numbers of neurons from the granular a subregion or the dysgranular RSC were also included.

### 2.6 Data analysis

#### 2.6.1 Firing Rate Maps

The T-maze was divided into 2.5 × 2.5 cm square pixels, and the firing rate of each neuron was determined by dividing the total number of spikes in each pixel by the time spent in the pixel. Spatial firing rate maps were smoothed by convolution with a 7×7 pixel Gaussian kernel with unity sum.

#### 2.6.2 Firing during the delay period

We determined whether RSC neurons exhibited differential firing during left and right delay period by examining firing patterns at the level of individual neurons and at the population level. In order to determine whether individual neurons carried information about the trial type (left or right) during the delay period, we computed the average difference in firing rates (z scores) for even and odd trials of the same trial type (left VS left and right VS right). This served as a baseline measure of variability in firing across trials of the same type. We then compared that to the difference in firing rates for trials of the opposite trial type (left VS right) for each neuron. This approach benchmarks any left-right firing rate differences against the baseline variability for each neuron. We also computed a t-score (Student’s *t*) comparing the firing rates for left and right trials for each neuron. This serves as a measure that reflects the trial-by-trial reliability of firing rate differences for left and right delay periods.

To assess left-right differential firing at the population level, we combined the neurons from all rats and recording sessions into a single population and used a minimum distance classifier to predict the trial-type based solely on neuronal firing patterns during the delay period. Population firing rate vectors for every 200 msec time bin during the 30 sec delay period were constructed using *z*-scored firing rates to control for differences in baseline firing, which can make population measures overly sensitive to a few neurons with high firing rates. We then computed the Euclidean distance between each of these population vectors and the mean firing rate vector for the left trial-type and the right trial-type and assigned the sample vector to the closest match (smallest distance). This analysis asks how well the firing rates match the average for each trial-type at each timepoint during the delay period. Decoding success reflects how reliably the firing rate at each of these timepoints differentiates the two trial-types.

#### 2.6.3 Firing at the trial start

First, firing rate of each neuron was calculated in 200 msec bins, and the values were averaged across the correct trials for each trial type (left and right) for the entire duration of 30 sec delay. Then, for each trial type, if the maximal firing during the 2 seconds before the delay end was greater than 3 standard deviations above the mean firing of entire delay period, it was classified as having a trial start response. Another criteria was imposed to eliminate the neurons whose elevated firing near delay end was solely due to the movement variables like running speed or acceleration. We calculated the correlation between firing rate and speed and the correlation between firing rate and acceleration for the entire session. Any neuron with a correlation (Pearson’s *r*) greater than 0.2 for either speed or acceleration was excluded.

## 3. Results

RSC lesions caused a significant impairment in the delayed alternation task. A mixed effects ANOVA comparing control and lesion group performance over the three delay lengths revealed a significant main effect of delay duration (F(1,47) = 26.75, p < 0.001) and a significant main effect of the lesion condition (F(1,47) = 13.05, p < 0.001; Fig. 1B). The delay by lesion interaction just failed to achieve significance (F(1,47) = 3.71, p = 0.06), suggesting a trend toward a larger lesion impairment at greater delay lengths. A planned comparison of the two lesion conditions at the 30 sec delay found a significant lesion-induced impairment (t(15) = 2.93, p < 0.01), confirming that this delay duration was suitable for the subsequent recording study. The size of the lesions varied considerably (39.5 – 72.8% of the RSC, Fig. 1C), and the impairment at the 30 sec delay was correlated with lesion size (r = −0.78, p < 0.05; Fig. 1C). The extent of RSC lesions is shown in Fig. 1D.

Previous studies have documented the spatial firing properties of RSC neurons as rats traversed the maze during continuous alternation (Miller et al., 2019; Subramanian et al., 2024) and similar spatial firing patterns were observed here (see examples in Fig. 2A). In the present study, we focused our analyses primarily on the firing patterns occurring during the delay period, which have not been examined previously. We first sought to determine whether RSC firing patterns differed before left and right trials in a manner that could support memory during the delay. We found that many RSC neurons fired at reliably different rates for left and right trials throughout the delay (Fig. 2A). In order to examine the degree to which each of the recorded neurons participated in such differentiation, we computed a between-trial type difference score by subtracting the z-scored firing rates for left and right trials, and then compared that to a within-trial type difference score computed by subtracting firing rates for trials of the same type (i.e. left VS left and right VS right). This is a particularly conservative approach because we not only ask whether there is a difference between left and right trials, but we also ask whether this difference is greater than the average variation across trials of the same type. We found that the difference between trial types was significantly greater than for trials of the same type when the rats performed correctly (t(143) = 4.08, p<0.001), but this differential firing was absent on trials when the rat made an error (t(104) = 0.33, p = 0.73).

**Figure 2.**
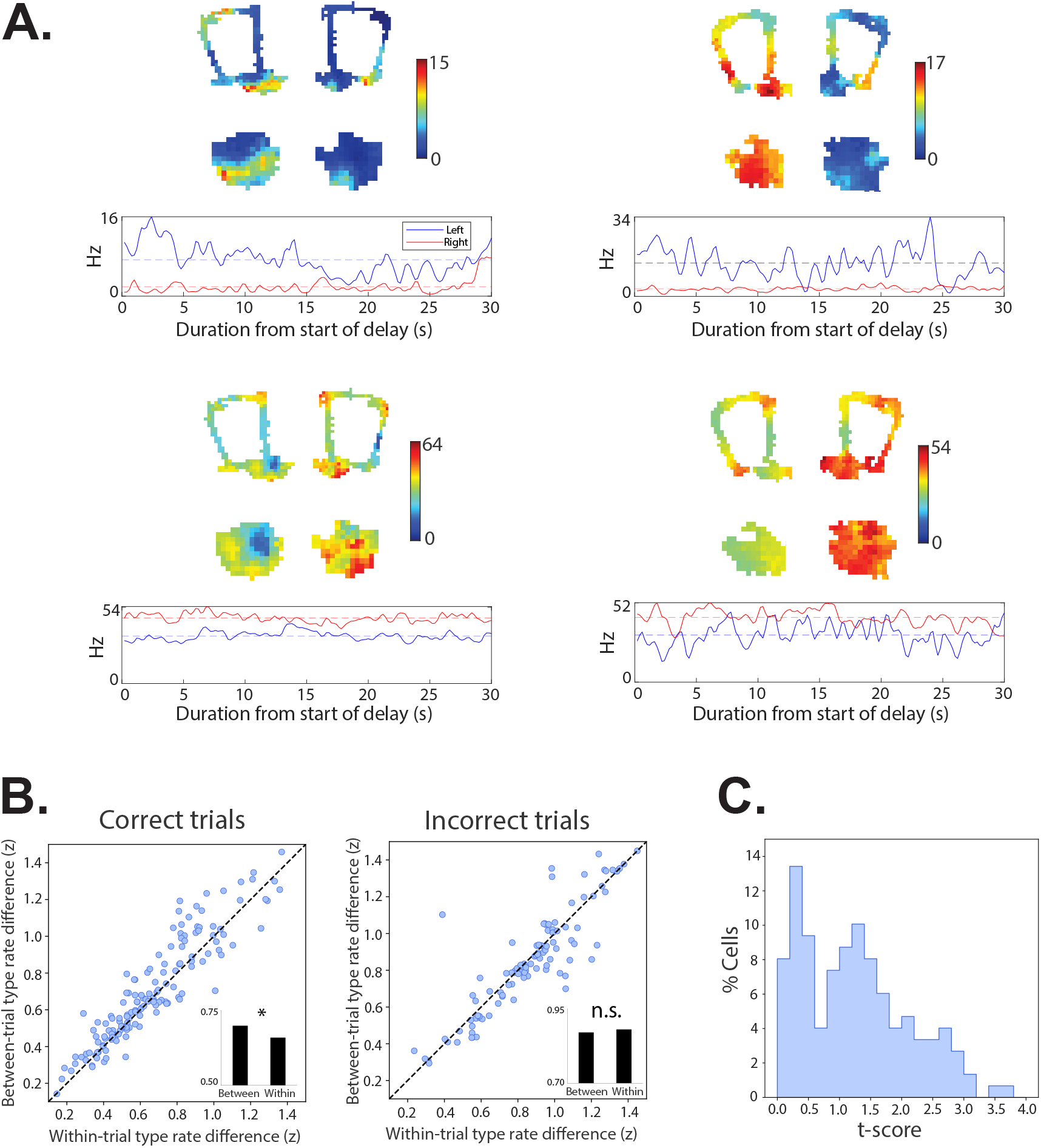
RSC neurons fire differentially during left and right delay periods. A) Four example neurons with distinct firing rates during the left and right delay periods are shown. Spatial firing rate maps are shown for the full session, with left and right trials plotted separately, illustrating the trials (above), and enlarged images of the delay area (below). The line plots show the average firing rate during the 30 sec delay across all left (blue) and right (red) trials. B) The difference in z-scored firing rates for left and right trials are plotted against the difference between even and odd trials of the same type (left or right) for each neuron (dots). The values for the correct trials are plotted on the left and the incorrect trials on the right. For correct trials, note that majority of the dots are above the unity line, indicating that these neurons differentiated right and left trials. In contrast, on trials where the rat subsequently made an error, fewer neurons differentiated the two trial types (dots were close to, or below the unity line). The bar graph insets show the average of the within- and between trial-type difference scores. C) The distribution of t-scores comparing the trial-by-trial firing rates during the left and right delay periods for each neuron is shown. Note the bimodal distribution, with a group of neurons with t-scores above 1.0 representing a population of neurons that fire at reliably different rates during the delay periods before the left and right trials.

The above-described firing rate differences could have been driven by extreme firing on a few trials, rather than reliably different firing across most trials. We examined this by computed a t-score (Student’s *t*) comparing the trial-by-trial firing rates from the delay periods preceding the left and right trials. The distribution of these values was bimodal (Fig. 2C), with a clear grouping of neurons with large t-scores suggesting that a large subset of the neurons reliably differentiate the left and right trial types. To examine this at the population level, we used a minimum distance classifier to predict the trial type (left or right) solely from the population firing patterns (see Methods). Consistent with the analysis of individual neurons, we found that we could correctly classify the left and right delay period from the population firing patterns 78.73% of the time on correctly performed trials, but classification accuracy was reduced to 61.73% on error trials.

Our previous study using a continuous (non-delayed) version of this T-maze task (Miller et al., 2019, Subramanian et al., 2024) found that RSC neuronal firing patterns differentiate left and right trials as the rats traverse the stem of the maze as well as the left and right reward locations. In the present study, we examined stem- and reward-related firing using the same strategy we employed for the delay period. For each neuron, we compared between- and within-trial type difference scores for firing rates during left and right stem traversals and during the two seconds after arrival at the left and right reward locations. We found that during stem traversal on correctly performed trials, the between-trial type difference was significantly greater than the within-trial type difference (t(120) = 2.82, p<0.01), indicating that the neurons reliably differentiated the stem traversals during left and right trials. These differential firing patterns were reduced on trials when the rat made an error (t(87) = 1.75, p = 0.08). Using the same analysis approach, we found that RSC neurons clearly differentiated the left and right reward locations (t(143) = 5.13, p<0.001). Analysis of population firing patterns using the minimum distance classifier as described above found that we could correctly classify the left and right trials from firing on the stem 75.64% of the time, but only 59.83% of the time on error trials. Population firing patterns differentiated the two reward locations with near-perfect accuracy (99%) on correctly performed trials. Reward-related firing cannot be assessed on incorrect trials since the rat does not receive a reward after an error.

Notably, there was no clear relationship between differential firing during the delay and differential firing on the stem or at the reward locations. To examine this, we correlated the firing rate difference scores for left and right trials for the delay period with the difference scores for left and right stem traversals and reward-related firing. If the same neurons that fired differently for the left and right delay periods were the same neurons that fired differently during stem traversals or at the reward, we expect these correlations to be high. However, correlations were low and non-significant for both comparisons (Fig. 3, stem: r = 0.01, p = 0.91; reward: r = 0.06, p = 0.53).

**Figure 3.**
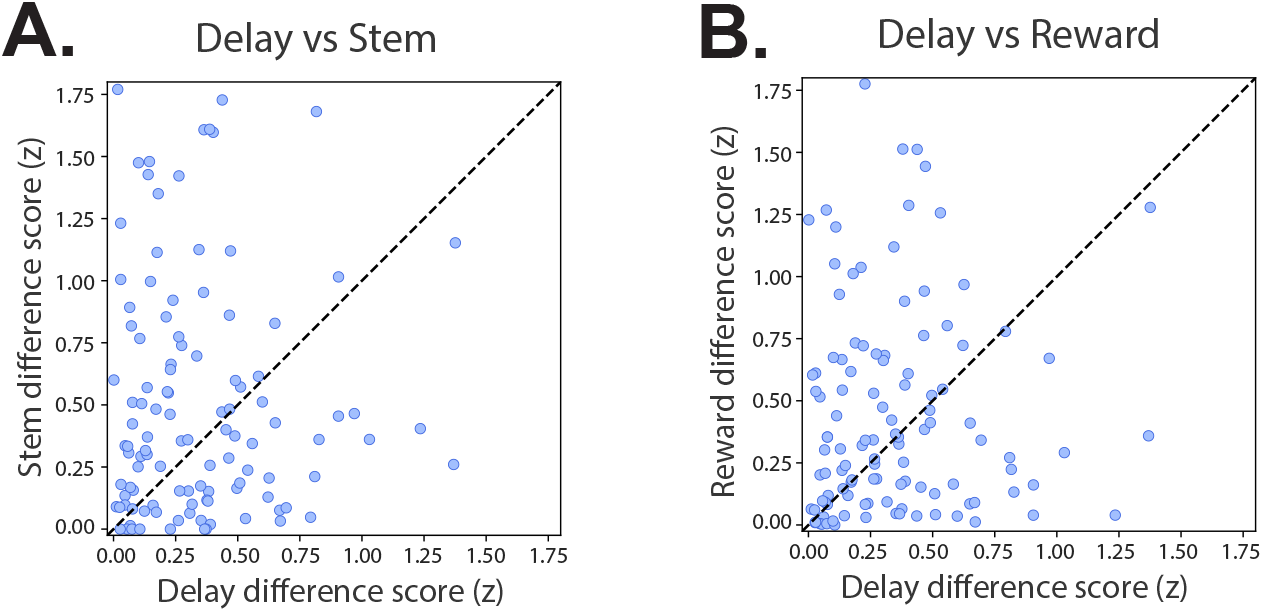
Differential firing during the delay period is not correlated with differential firing on the stem or at the reward locations. A) The degree to which neurons differentiated the left and right delay periods (z-scored difference scores) was not correlated with the degree of differentiation during stem traversal. B) Differential firing during the delay was also not correlated with differential firing at the reward locations.

We observed that many RSC neurons exhibited a sudden spike in firing rate leading up to the start of the trial when the block was removed (Fig. 4). This spike in firing could serve as a ‘go signal’ used to initiate the behavioral response, so we examined the prevalence of this signal in our data set. We binned the average firing rate into 200 msec bins for the entire duration of 30 sec delay, and then classified the neurons as having a putative go signal if their maximum firing rate during the final 2 sec of the delay period was at least 3 standard deviations above the mean of the entire delay period. The firing rate of some RSC neurons is strongly correlated with running speed or acceleration (Subramanian et al., 2024), which could cause an increase in firing at the start of the trial solely as a consequence of the rat’s locomotor behavior. After excluding these movement-sensitive neurons (see Methods), we found that 29% (42/144) of RSC neurons met our criteria for having a trial start signal (Fig. 4A-B). Of these, 20 had a trial start signal for both left and right trials, while 22 had trial start firing for only one type of trial.

**Figure 4.**
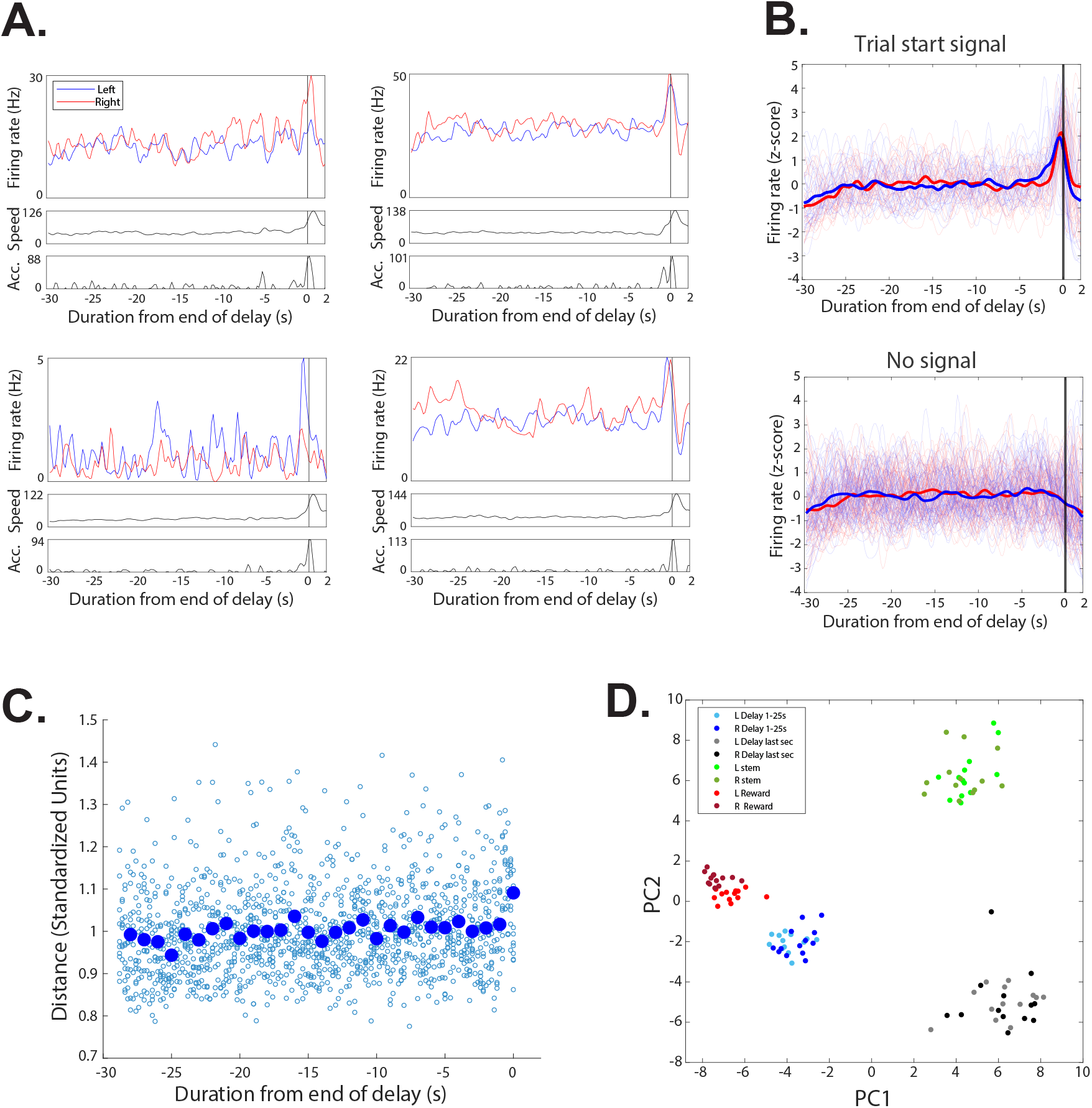
Many RSC neurons exhibited a spike in firing rate just before the start of the trial run. A) Four example neurons with a sudden increase in firing before the start of the trial (vertical line). Average firing rates are shown separately for left (blue) and right (red) trials for the full 30-sec delay period and for 2 sec into the trial run. Below, the rat’s average speed (cm/sec) and acceleration (cm/sec^2^) are plotted. About half of the neurons exhibited greater trial start firing for either left or right trials (left plots). B) The firing rates of all neurons that met our criteria for trial start firing (top) are overlayed with the mean of each trial type (bold line) along with all the neurons that did not (bottom). C) The change in population firing patterns across the delay period is shown. Small circles illustrate the Euclidean distance between the first second of the delay period and each successive 200 msec time bin. The large circles show the mean for each 1-sec interval. Note that the activity is relatively stable for most of the delay period (dots form a flat line). However, the population firing pattern suddenly changes during the last second of the delay (elevated distance value), reflecting the shift in firing before the start of the trials. D) The first and second principal components are shown for the population firing patterns during the first 25 sec of the delay period, the last sec of the delay, the stem traversal, and arrival at the reward location, with left and right trials shown in light and dark colors. Note that population firing patterns are distinct for each of these trial epochs, including the last second of the delay period.

We confirmed that this trial start response was detectable within the broader population of all RSC neurons in two ways. First, we generated population vectors including all of the recorded neurons for each 200 msec time bin during the delay period and we measured the Euclidean distance between each of these population vectors and the vector for the first second of the delay. This serves as a measure of how much the population firing patterns changed during the delay period, relative to the start of the delay period (Fig. 4C). We found that the population firing patterns were relatively stable until the final second before the start of the trial, when the distance became significantly greater than the average of the previous seconds (t(49) = 5.54, p<0.001). This suggests that the population firing patterns exhibited a sudden shift immediately before the start of the trial, presumably due to the presence of the trial start responses within the population. We also subjected the population firing patterns to a principal components analysis, which included population vectors constructed from firing during stem traversal, at the reward, during the first 25 seconds of delay, and final second of delay. The first two components of the PCA clearly differentiated the firing patterns during each of these epochs, including the time of the trial start response as a distinct activity state of the RSC (Fig. 4D).

## 4. Discussion

RSC lesions significantly impaired delayed alternation performance, consistent with previous reports of impaired spatial working memory in subjects with RSC lesions (Keene & Bucci, 2009; Vann & Aggleton, 2004). We also recorded RSC neuronal activity and found firing patterns similar to those reported previously in a continuous version of the task. For example, we confirmed that RSC neurons fire differentially on the stem of the maze for left and right trials (Miller et al., 2019) and they reliably differentiate the two reward locations (Miller et al., 2019; Smith et al., 2012). We also report two new findings: First, RSC neurons exhibit reliably different firing patterns during the delay periods preceding left and right trials, consistent with a working memory signal. Second, we discovered that many RSC neurons exhibit a large spike in firing rate leading up to the start of the trial.

Behavioral performance during the 0-sec delay condition was very good (~90% correct), which is similar to our previous study of continuous alternation (Miller et al., 2019). For this condition, we placed a block at the entrance to the stem, just as we did in the delayed trials, but we removed the block as soon as the rat arrived at the start position. The strong performance for this condition suggests that the mere presence of the block did not distract the rats or otherwise disrupt performance. Performance declined significantly with increasing delay duration, suggesting that the rats rely on a working memory strategy rather than simply learning to traverse the maze in a continuous figure eight pattern, which is possible in continuous versions of the task. If performance had depended critically on running along a specific figure eight trajectory, then even the brief disruption of smooth running in the 0-sec delay should impair performance and increasing the delay duration would not be expected to produce a greater disruption of trajectory coding.

Rats with RSC lesions were significantly impaired in this task, particularly in the 30-sec delay condition. The severity of the impairment was closely correlated with the extent of RSC tissue damage. Very large lesions produced a severe deficit, with performance near chance levels, but smaller lesions that destroyed less than half of the RSC tissue produced variable performance that was similar to controls (see Fig. 1C). The lesions targeted all subdivisions and the full rostro-caudal extent of the RSC. The largest lesions necessarily included more of the rostral and caudal tissue and more extensive damage to the granular a subregion (Rga). However, the range of lesion placement did not allow systematic examination of the role of individual subregions. Our previous studies of RSC neuronal firing patterns (Subramanian et al., 2024) suggest a distributed coding scheme with little evidence of specialization along the rostro-caudal extent of the RSC (but see Pothuizen et al., 2009; Trask et al., 2021; Van Groen et al., 2004).

RSC neurons exhibited well-differentiated firing during the delay period before left and right trials. Although the degree to which individual neurons differentiated the trial types varied, some neurons fired at different rates continuously throughout the delay period (see examples in Fig. 1A). These neurons are notably similar to our recent findings on RSC differentiation of environmental contexts (Miller et al., 2021), in which many RSC neurons exhibited reliably different firing rates throughout the twelve-minute foraging trials in black and white boxes. In that study, the differential firing rates could have been driven by the different visual input in the black and white boxes. In the present study however, visual input did not differ systematically between the left and right delay periods, indicating that RSC neurons can differentiate conditions based on internally generated memory demands and need not be driven by external stimuli (also see Sun et al., 2021). These left-right firing rate differences were significant, even when compared to a conservative baseline of variation across trials of the same type and importantly, they were absent during delay periods preceding an error.

In combination with the lesion-induced impairment, these data strongly suggest that these differential firing patterns play a critical role in the memory representations needed to perform this task. Previous studies have found that RSC neurons encode information about goal locations as well as trajectories through space (e.g. Alexander and Nitz, 2015, 2017; Miller et al., 2019), so the delay-related firing seen here could be linked to either the specific goal location or the route to the goal. However, our data indicate that the delay-related firing was not correlated with reward location firing or firing during stem traversal. In other words, neurons with greater firing during the delay period before right (or left) trials were not necessarily the same neurons that had greater firing on right (or left) stem traversals or reward-related firing. Previous work has shown that RSC neurons typically encode multiple task variables and key task features are encoded by overlapping populations of neurons (e.g. Subramanian et al., 2024, Vedder et al., 2017). Our current results are consistent with this insofar as different but overlapping populations of RSC neurons encoded right and left trial type information during the delay period, stem traversal and at the reward locations.

We also found that many RSC neurons ramped up their firing rate just before the rats began their run down the stem of the maze. The functional role of these firing patterns is not known, and there are several plausible explanations. We excluded neurons with firing rates that were correlated with running speed and acceleration, so it is not likely that these firing bursts are a simple consequence of locomotor activity. The observation that approximately half of the neurons ramped up their firing selectively on left or right trials even though the locomotor behavior did not differ appreciably further suggests that the firing is not a consequence of the locomotor activity. Nevertheless, the firing invariably occurred near the time when the rats began the trial run, making it impossible to fully disentangle the neuronal firing from the behavioral response. Consistent with this idea, a previous study found that RSC neurons ramp up their firing to the time of a locomotor avoidance response (Kubota et al., 1996), and these authors suggested that this might represent a ‘go’ signal which could be transmitted to the striatum to initiate the behavioral response.

Previously, we found that hippocampal neurons exhibit a similar spike in firing rate at the start of the trials in an odor discrimination task (Bulkin et al., 2019). Specifically, a large spike in firing occurred when a barrier was removed to allow rats to leave the intertrial waiting area and enter the area where the odor cues were presented. We found that large scale population firing patterns shifted from a ‘waiting’ state to a ‘trial performance’ state and we speculated that the firing rate spike could serve to drive hippocampal representations from one state to the other. The hippocampus is interconnected with the RSC (Sugar et al., 2011; Wyss & Van Groen, 1992). If a similar spike in hippocampal firing occurs at the start of trials in the delayed alternation task, this could drive the spike in firing in the RSC, as well as the apparent shift in population firing patterns between the delay period and the stem run suggested by our principal components analysis (Fig. 4D).

Overall, these results support an RSC role in spatial working memory and the differential firing patterns during the delay period represent a plausible mechanism for maintaining the working memory representations needed for successful task performance. Additionally, the spike in firing at the start of the trial runs may trigger the execution of the motor programs supporting the ‘go right’ and ‘go left’ behavioral responses and shift the population representations into the appropriate state for trial performance. Much of the literature on the RSC has focused on how firing patterns reflect information relevant to ongoing experience, such as the subject’s current spatial location and heading direction. The present results, in combination with previous findings that RSC neurons simulate future goal locations (Miller et al., 2019), indicate that RSC activity is also concerned with future behavioral choices.

## Acknowledgements

This work was supported by MH083809 to D. Smith.

